# Lipid-laden endothelial cells exhibit a transcriptomic signature linked to blood-brain barrier dysfunction, metabolic reprogramming and increased inflammation in the aging brain

**DOI:** 10.1101/2025.08.22.671845

**Authors:** Sarah Otu-Boakye, Duraipandy Natarajan, Bhuvana Plakkot, Ilakiya Raghavendiran, Tamas Kiss, Madhan Subramanian, Priya Balasubramanian

## Abstract

Dysregulation in lipid metabolism is increasingly recognized as a key contributor to age-related diseases, including neurodegeneration and cerebrovascular dysfunction. While prior studies have largely focused on glial cells, the impact of lipid dysregulation on brain endothelial aging remains poorly understood. In this study, we conducted a secondary analysis of single-cell transcriptomic data from young and aged mouse brains, with a specific focus on endothelial cells (ECs). Our analyses revealed that aging promotes lipid droplet accumulation in brain ECs. These lipid-laden brain ECs exhibit a transcriptomic signature indicative of impaired blood-brain barrier function, increased cellular senescence, and inflammation in aging. Furthermore, lipid accumulation is associated with an altered metabolic phenotype characterized by increased fatty acid oxidation and decreased glycolysis, and impaired mitochondrial electron transport chain activity in the ECs of the aging brain. We have also validated lipid accumulation in aged ECs *in vivo*. Collectively, our findings indicate that lipid accumulation drives structural, functional, and metabolic impairments in the brain ECs, likely contributing to cerebrovascular aging. Understanding the mechanisms underlying lipid accumulation-induced endothelial dysfunction may offer novel therapeutic strategies for mitigating microvascular dysfunction and cognitive decline in aging.

## Introduction

Lipid droplets (LDs) are intracellular organelles that serve as key regulators of lipid storage and mobilization in response to the metabolic demands of the cell. Under conditions of excess energy availability, cells esterify free fatty acids (FAs) into triglycerides (TGs) and sequester them within LDs to prevent lipotoxicity. Conversely, during periods of energy deprivation or increased metabolic demand, TGs stored in LDs undergo hydrolysis by lipases such as adipose triglyceride lipase (ATGL), releasing FAs and other lipid metabolites that can be utilized for energy, membrane biosynthesis, and signal transduction^1^. While LDs are present in most cell types under physiological conditions, excessive LD accumulation is often associated with pathological states. Most studies investigating abnormal LD accumulation have focused on peripheral tissues, particularly hepatocytes in the context of metabolic-associated steatohepatitis (MASH) and adipocytes in obesity. However, growing evidence suggests that dysregulated LD metabolism also plays a critical role in neurodegenerative diseases^2-7^.

In the central nervous system (CNS), pathological LD accumulation has been primarily observed in microglia and astrocytes, where it contributes to neuroinflammation and neurodegeneration in aging and Alzheimer’s disease (AD) models^2,4^. LD accumulating microglia, termed as LDAM, in the aging brain exhibit a pro-inflammatory phenotype with excessive reactive oxygen species (ROS) and altered phagocytic activity^2^. Similarly, astrocytic LD accumulation has been linked to senescence^3^ and microglial activation and inflammation in obesity^8^. In addition, ectopic LD accumulation in the ependymal cells is linked with attenuated neural stem cell proliferation and cognitive impairment in AD^9^. Conversely, studies also highlight a neuroprotective role for glial LD accumulation. ATGL inhibition-induced LD accumulation in microglia has been shown to reduce infarct volume and improve neurocognitive outcomes in acute ischemic stroke^10^. Likewise, astrocytic LD accumulation protects neurons by buffering stress-induced lipid overload and preventing lipotoxicity^11^. These findings underscore that the effects of LDs are highly context- and cell-type-specific, exerting both detrimental and protective roles depending on the disease state and cellular environment.

Dysregulated lipid metabolism is emerging as a major player in brain aging^12-16^; however, these studies have primarily focused on glial cells and neurons. It is not clear whether aging also induces lipid dysregulation in the vascular cells, specifically brain endothelial cells (ECs), which play a critical role in regulating blood flow responses and maintaining blood-brain barrier (BBB) integrity. Given that microvascular dysfunction is a key driver of vascular cognitive impairment (VCI) and neurodegeneration, understanding whether age-related lipid dysregulation in ECs contributes to cerebrovascular aging and cognitive decline is of significant therapeutic interest. Previous analysis in single-cell data sets has highlighted lipid biosynthesis and metabolism as one of the significantly altered pathways in brain ECs with aging^17^. However, a comprehensive analysis of how aging affects lipid metabolism in brain ECs and how these changes impact the function and phenotype of brain ECs remains lacking. Understanding these knowledge gaps could uncover novel therapeutic targets aimed at restoring lipid homeostasis in brain ECs to mitigate age-related endothelial dysfunction and cognitive impairment. To address this, we re-analyzed a publicly available single-cell RNA sequencing dataset published by Ximerakis et al.^17^, in which transcriptomic profiling of ∼50,000 brain cells, including ECs, was performed in young and aged mice. We specifically focused on transcriptional changes in pathways related to lipid synthesis, oxidation, and turnover, and investigated their relationship to known age-related pathological changes in brain ECs, including BBB dysfunction, inflammation, and cellular senescence.

## Methods

### Pre-processing of single-cell data

Single-cell transcriptomic data were obtained from the study “Single-cell transcriptomic profiling of the aging mouse brain” (Ximerakis et al., 2019; https://doi.org/10.1038/s41593-019-0491-3), available via the Broad Institute’s Aging Mouse Brain Portal. The dataset contained a total of 37069 cells; no additional quality control (QC) steps were applied beyond those described in the original study.

A Seurat object (v5.1.0, R v4.4.1) was created from the raw count matrix and metadata. Initial normalization was performed using the LogNormalize method with a scale factor of 10,000. Highly variable features (n = 2000) were identified using the VST method. The data were scaled, and principal component analysis (PCA) was conducted on the variable features. The top 20 principal components (PCs) were used for further analysis. Graph-based clustering was performed using the FindNeighbors and FindClusters functions with a resolution of 0.05, resulting in 25 transcriptionally distinct clusters. Cells were annotated based on canonical marker expression. Cell clusters were visualized using Uniform Manifold Approximation and Projection (UMAP), a non-linear dimensionality reduction method.

### Gene set enrichment analysis

Gene set activity scores were calculated using the AUCell R package (v1.26.0). Custom gene sets were filtered to include only genes present in the dataset. AUC scores were computed for each gene set and cell, and the resulting values were added to the Seurat object metadata for downstream visualization and analysis^18^. The list of genes for each set analyzed is provided in Table 1. Senescent cells were classified according to a combined senescence signature and Cdkn2a expression criterion^19^. Cells were defined as senescent if they were either Cdkn2a_pos or belonged to the “top” senescence score group. Cells were defined as non-senescent if they were Cdkn2a_neg or belonged to the “bottom” senescence score group.

**Table 1.**
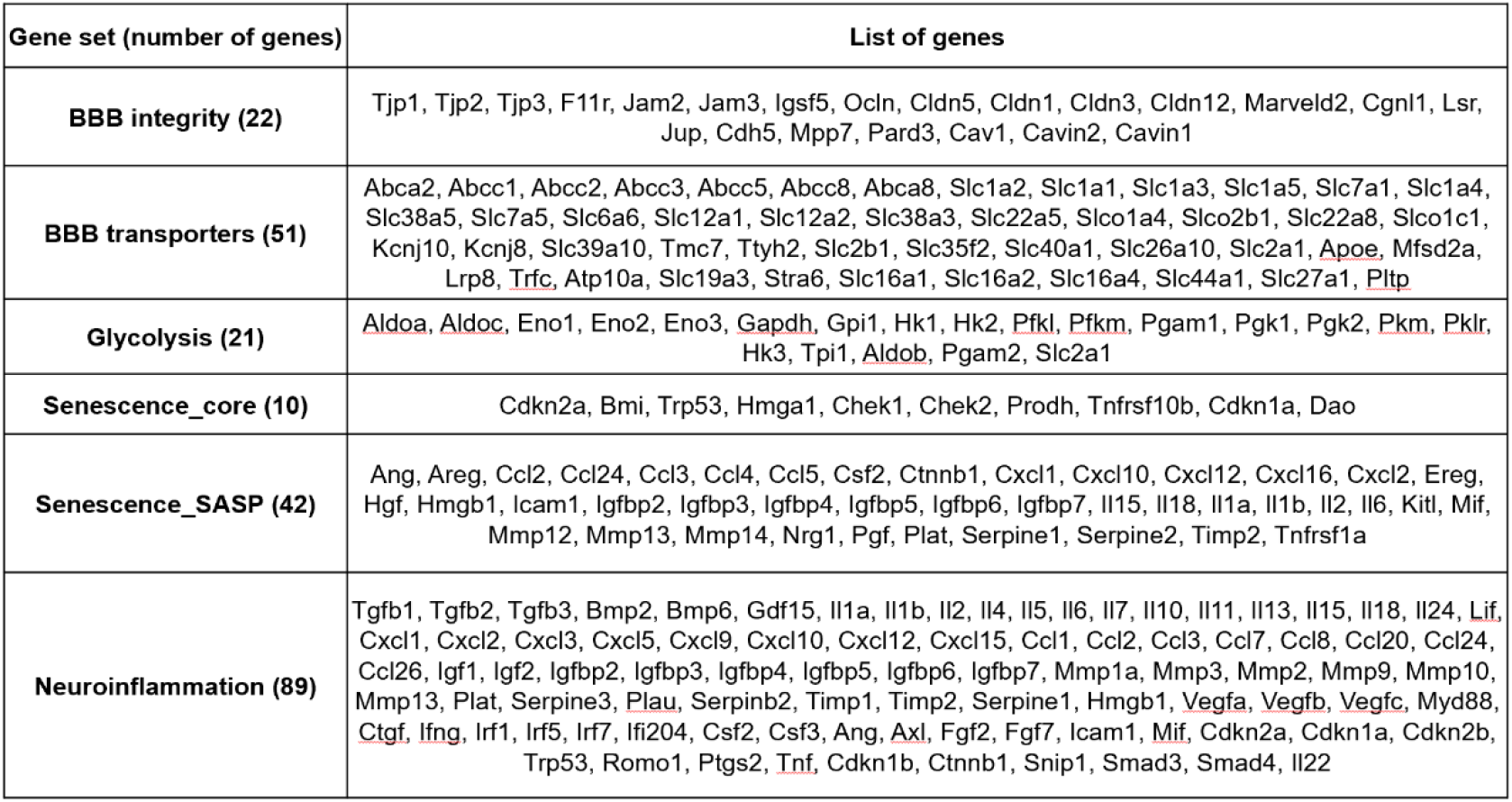
List of genes used in the gene set enrichment analysis.

### Endothelial cell isolation from mouse brain

Brains from young (3 months old) and aged (24 months old) male and female mice were processed using the Adult Brain Dissociation Kit (Miltenyi Biotec, #130-107-677). Following transcardial perfusion with 1X phosphate-buffered saline (PBS), brains were dissected, minced, and enzymatically dissociated according to the manufacturer’s protocol. Tissue homogenates were filtered sequentially through 100 µm and 30 µm MACS SmartStrainers and centrifuged to obtain single-cell suspensions. Myelin and cellular debris were removed by density gradient centrifugation (centrifuged at 4°C and 3000g for 10 minutes with full acceleration and full brake), followed by red blood cell lysis using the 1X red blood cell removal solution. For endothelial cell enrichment, single-cell suspensions were incubated with CD31 microbeads (Miltenyi Biotec, #130-097-418, 10ul per sample) for 15 min at 4°C, washed, and applied to LS MACS columns (Miltenyi Biotec, #130-042-401) placed in a magnetic field. After washing, CD31^+^ endothelial cells were eluted with MACS buffer and collected by centrifugation (300g for 10 minutes). The enriched endothelial cell fraction was resuspended in appropriate 1X lysis buffer for downstream protein analyses.

### Capillary-based immunoassay for protein analysis

Isolated endothelial cells from young and aged mice were lysed in 1X cell lysis buffer (Cell Signaling Technology, Danvers, MA, USA) supplemented with Halt protease and phosphatase inhibitor cocktail (Thermo Fisher Scientific, #PI78440). The lysates were clarified by centrifugation at 16,000 × g for 10 min at 4 °C. The resulting supernatant was collected, and protein concentration was quantified using the Pierce BCA Protein Assay Kit (Thermo Fisher Scientific, #23227).

Automated western blot analysis was carried out using the Jess capillary-based immunoassay platform (Biotechne, ProteinSimple) with a 12–230 kDa separation range and the Protein Normalization (PN) module, processed through Compass for SW Software (version 6.2.0). Protein samples were diluted in 0.1X sample buffer and loaded at an optimized concentration of 1 µg/µL for ATGL, PLIN2, DGAT1 and GAPDH; 0.5 µg/µL for MCAD. The following antibodies were used: rabbit anti-ATGL antibody (Cell Signaling Technology #2138S) at 1:50 dilution, rabbit anti-PLIN2 antibody (Proteintech #15294-1-AP) at 1:50 dilution, goat anti-DGAT1 antibody (Novus #NB100-57086) at 1:50 dilution, mouse anti-GAPDH antibody (Cell Signaling Technology #97166S) at 1:50 dilution and mouse anti-MCAD antibody (Santa Cruz Biotechnology #sc365108) at 1:10 dilution. Peak areas corresponding to ∼54 kDa for ATGL, ∼45-48 kDA for PLIN2, and ∼45 kDa for MCAD were quantified via dropped line peak integration and normalized to total protein levels using the PN module in Compass for SW Software. DGAT1 (∼55 kDa) protein levels were normalized to GAPDH (∼40 kDa) protein levels.

### Statistical analysis

Statistical analyses were performed using Graph pad prism 9.3.1 (GraphPad Software, San Diego, CA, USA) and the data are expressed as mean±SEM. Data were analyzed by two-tailed, unpaired student’s t-test and p<0.05 were considered statistically significant.

## Results

### Brain ECs exhibit decreased lipid turnover and increased lipid accumulation in aging

To assess the impact of aging on lipid metabolism in brain ECs, we conducted a secondary analysis of a publicly available single-cell RNA sequencing dataset published by Ximerakis et al.^17^. This dataset provides a comprehensive transcriptomic profile of approximately 37000 brain cells capturing all brain cell types from young (2-3 months) and aged (21-23 months) brains. Louvain clustering analysis identified 25 different cell clusters, including the endothelial cell population **(Fig. 1A, dotted red box)**, based on the expression of classical endothelial-specific markers such as *Cldn5* and *Slco1a4*. This analysis enabled us to investigate the expression of lipid metabolism-related genes across various brain cell types at single-cell resolution. Notably, brain endothelial cells expressed genes encoding LD associated proteins (perilipin 2 (*Plin2*) and perilipin 3 (*Plin3*)) **(Fig. 1B)**, lipid hydrolyzing enzymes (*Pnpla2*, which encodes ATGL, and *Ddhd2*) **(Fig. 1C)** and lipid synthesizing enzymes (*Acly* and *Dgat1*) **(Fig. 1D)**. While perilipins were most abundantly expressed in glial cells, particularly oligodendrocytes which depend on lipids for myelin synthesis **(Supp. Fig. 1)**, our analysis confirms that brain ECs also express key genes involved in lipid metabolism.

**Figure 1.**
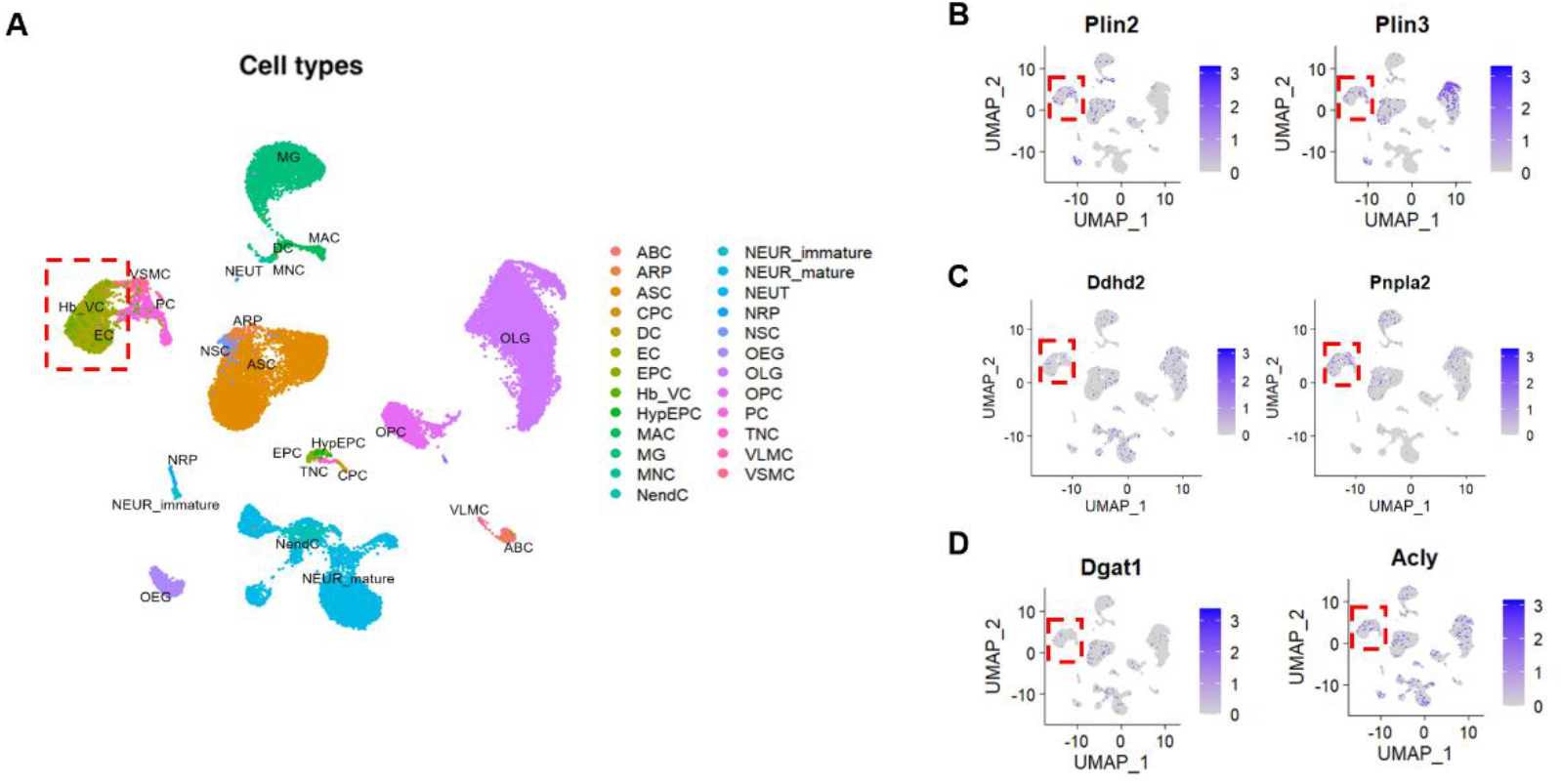
Louvain clustering analysis for the identification of cell types in the brain. **A.**Identification of 25 different cell clusters based on their transcriptional profile and differentially expressed genes as described in the original paper^17^. The cell clusters are color coded and annotated as follows: ABC-arachnoid barrier cells, ARP-Astrocyte-restricted precursors, ASC-Astrocytes, CPC-Choroid plexus epithelial cells, DC-Dendritic cells, EC-Endothelial cells, EPC-Ependymocytes, Hb-VC-Hemoglobin-expressing vascular cells, HypEPC-Hypendymal cells, MAC-Macrophages, MG-Microglia, MNC-Monocytes, NendC-Neuroendocrine cells, NEUR-immature-Immature neurons, NEUR-mature-Mature Neurons, NEUT-Neutrophils, NRP-Neuronal-restricted precursor, NSC-Neural stem cells, OEG-Olfactory ensheathing glia, OLG-Oligodendrocytes, OPC-Oligodendrocyte precursor cells, PC-Pericytes, TNC-Tanycytes, VLMC-Vascular and leptomeningeal cells and VSMC-Vascular smooth muscle cells. The dotted red box indicates the endothelial cell cluster. **B-D**. Two-dimensional UMAP plots for the lipid metabolism-related gene expression in the cell clusters. The dotted red box indicates the endothelial cell cluster. Relative expression values for each cell in each cluster are shown in a graded blue color.

To further investigate age-related changes in lipid metabolism, we assessed the expression of *Plin2* and *Pnpla2* in the brain ECs. Bubble plot analysis integrates information on both the gene expression level and the percentage of cells expressing the gene within each group. Using this approach, we found that aged ECs exhibited a concomitant upregulation of *Plin2* (LD marker) and downregulation of *Pnpla2* and *Ddhd2* (lipid hydrolytic enzymes) **(Fig. 2A)**. However, aging did not affect the percentage of cells expressing Plin2 as ∼16-17% of ECs expressed Plin2 in both young and aged brains **(Fig. 2B-C)**. Overall, our findings suggest that aging is associated with a transcriptomic signature indicative of reduced lipid turnover and increased lipid accumulation in brain ECs.

**Figure 2.**
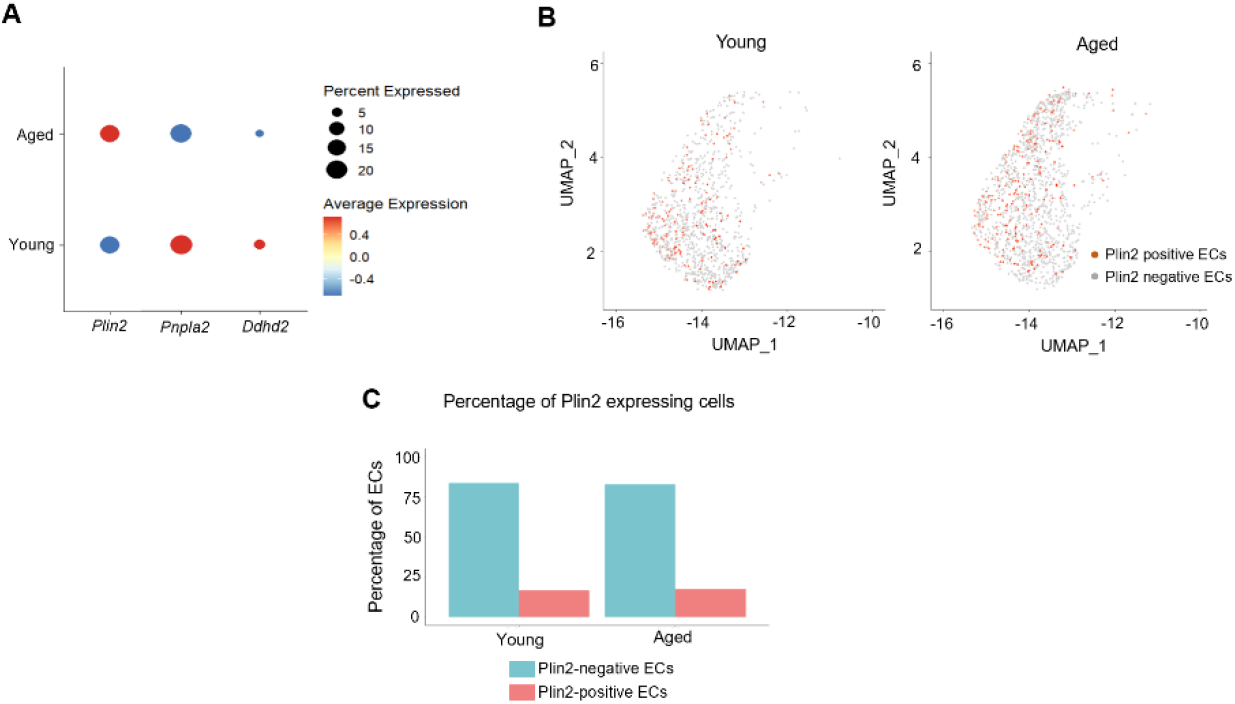
Age-related changes in endothelial lipid metabolism. **A.**Bubble plot analysis showing the expression level of *Plin2, Pnpla2*, and *Ddhd2* in young and aged ECs. Please note that the size of the bubble provides information on the percentage of cells expressing the gene, and the color indicates the directionality of the change in gene expression. **B**. Two-dimensional UMAP plots showing Plin2-expressing cells (dark orange colored cells) in the EC cell cluster in young and aged brains. C. Bar graphs showing the percentage of Plin2 expression cells in the young and aged brains.

### Lipid accumulation is associated with decreased expression of BBB-regulating genes in brain ECs with aging

Lipid accumulation, particularly in glial cells, has previously been shown to exert both beneficial and detrimental effects depending on the context and disease state. Therefore, it is important to understand how LD accumulation affects endothelial cell function and phenotype during aging. To investigate this, we performed gene set enrichment analysis focusing on genes that regulate the BBB, a critical function of ECs in the brain. Previous studies strongly support that age-related pathological changes in the ECs contribute to increased BBB permeability, leading to microglial activation and neuroinflammation. To assess how lipid accumulation affects BBB-related gene expression, we calculated a modified gene enrichment score for two gene sets: BBB integrity-related genes (n=22) and BBB transporter-encoding genes (n=51), specifically in Plin2-positive (LD accumulating) ECs from young and aged brains. This enrichment score incorporates both the expression level and the number of BBB-related genes expressed in each Plin2-positive cell. Our analysis revealed a substantial subpopulation of Plin2-positive ECs in the aged brain with reduced enrichment scores for both BBB integrity genes **(Fig. 3A)** and BBB transporter genes **(Fig. 3B)** compared to their counterparts in the young brain. We further performed a bubble-plot analysis for some of the key BBB-regulating genes to compare their expression between Plin2-positive versus Plin2-negative ECs in the aging brain. Consistently, LD-containing Plin2-positive ECs exhibited downregulation of critical genes involved in the maintenance of BBB integrity, like claudin5 (*Cldn5*), occludin (*Ocln*), tight junction protein 1 (*Tjp1*), CD31 (*Pecam1*), VE-cadherin (*Cdh5*), β-actin (*Actb*), and lipolysis-stimulated lipoprotein receptor (*Lsr*) **(dotted red boxes, Fig. 3C)** compared to Plin2-negative ECs. Together, these results suggest that lipid accumulation in ECs is associated with a transcriptomic profile indicative of impaired BBB function in aging.

**Figure 3.**
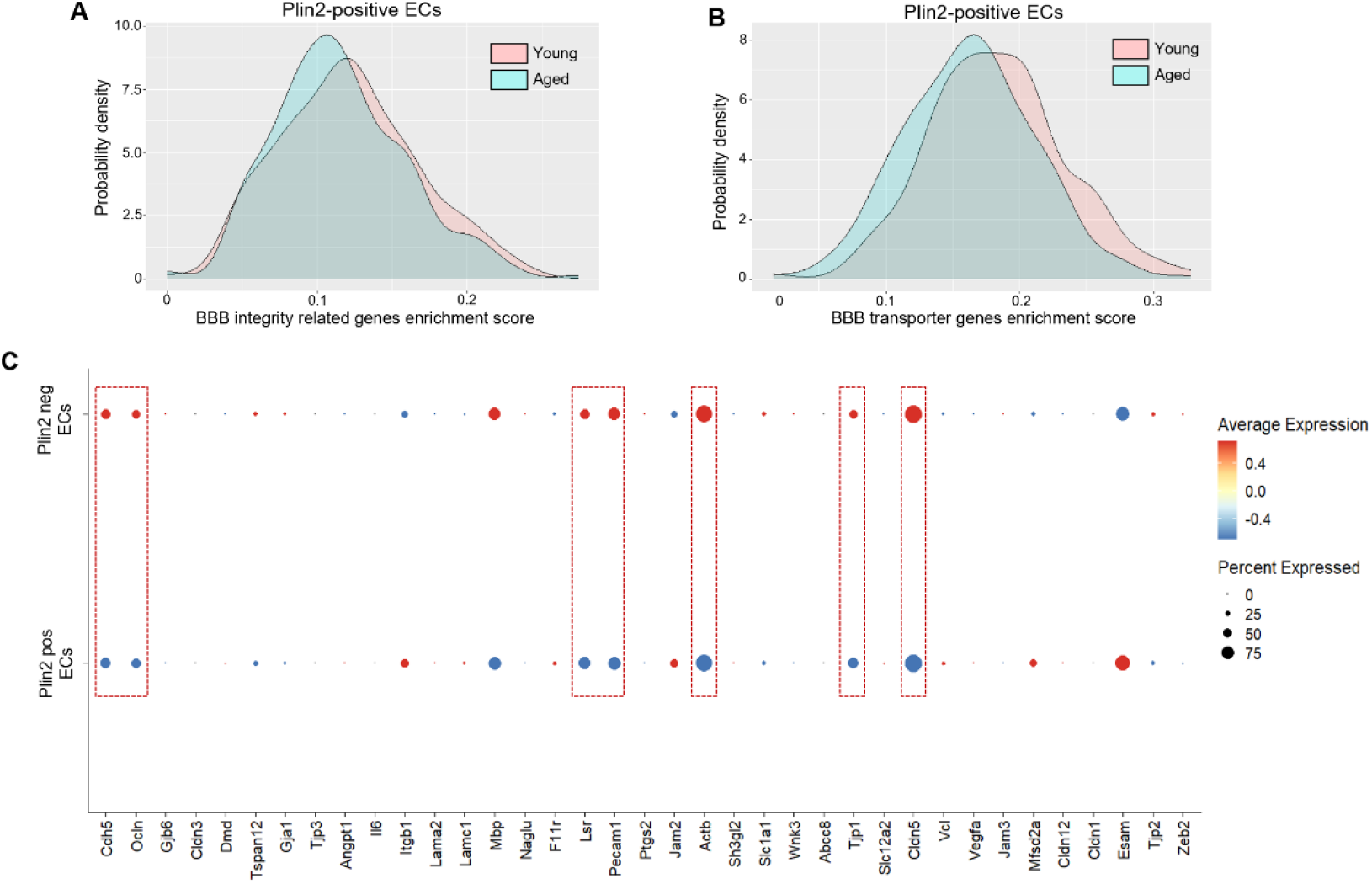
Changes in BBB-regulating genes in the Plin2-expressing brain ECs. **A-B.** Density plots for gene set enrichment scores calculated for BBB integrity and BBB transporter-related genes, respectively, in the Plin2-expressing ECs from young and aged brains. Please see the methods section for the list of genes used for this analysis. **C**. Dot plot analysis of some of the key genes regulating the integrity of the BBB between the Plin2-positive versus Plin2-negative ECs of the aging brain. Dotted red boxes indicate downregulation of tight and adherens junction proteins in the Plin2-positive ECs of the aging brain.

### Lipid accumulation correlates with increased lipid synthesis and oxidation, and impaired mitochondrial function in aged endothelial cells

Next, we assessed how lipid accumulation impacts energy metabolism in brain ECs. First, we investigated the expression levels of genes involved in lipid synthesis and oxidation. Several enzymes involved in fatty acid synthesis (*Acly, Dgat1, Dgat2, Acsl3, Acsl4* and *Acsl6*) and fatty acid oxidation (*Cpt1a, Cpt1c, Cpt2, Acsl1* and *Acadl*) were upregulated in the Plin2-positive compared to the Plin2-negative ECs in the aging brain (**Fig. 4A**). Since ECs primarily rely on glycolysis for energy^20^, we also evaluated the expression of glycolytic genes. Although we did not observe a uniform gene expression pattern among the glycolytic genes, some of the key rate-limiting enzymes, such as hexokinase (*Hk1* and *Hk2*), pyruvate kinase (*Pkm*), and the glucose transporter 1, GLUT1 (*Slc2a1*) were downregulated, indicating a potential metabolic shift from glucose to fatty acid utilization in the lipid-accumulating ECs (**Fig. 4B-C**). Interestingly, this phenotype appears to be endothelial-specific, as LDAMs, in contrast, exhibit increased expression of glycolytic genes (please see bubble plots and gene enrichment score density plots, **Fig. 5A-B)**, consistent with previous reports demonstrating a shift toward increased glycolysis as a hallmark of pathological microglial activation^21-23^. Overall, our data suggest that lipid accumulation drives metabolic reprogramming in a cell-type-specific manner.

**Figure 4.**
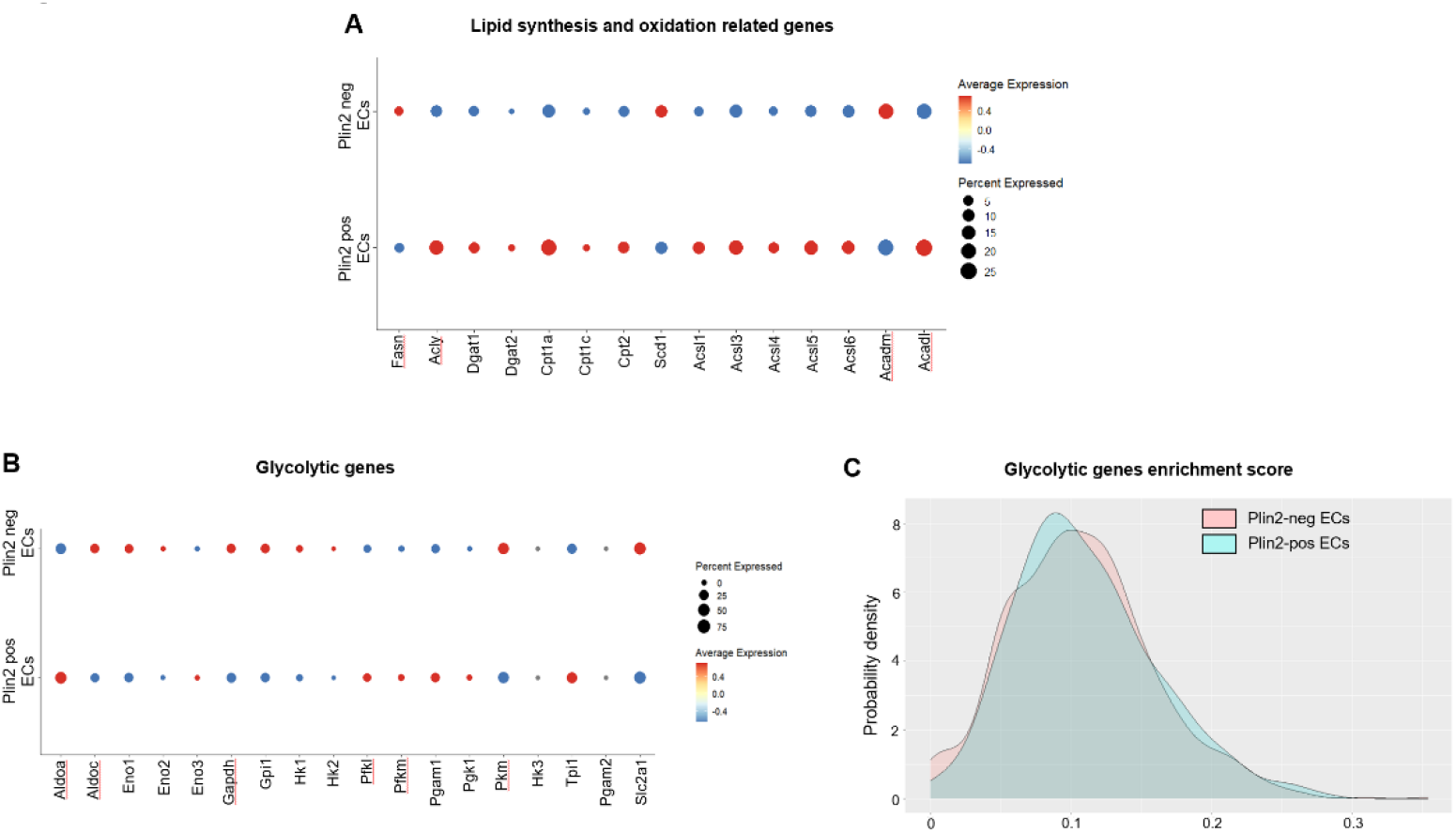
Changes in lipid synthesis and oxidation genes in the Plin2-expressing brain ECs. Dot plots showing the expression levels of genes involved in A) lipid synthesis and oxidation, and B) glycolytic pathway in Plin2 positive and negative ECs from the aged brains. C) Density plots for gene set enrichment scores for glycolytic genes in Plin2-positive and negative brain ECs from the aged brains.

**Figure 5.**
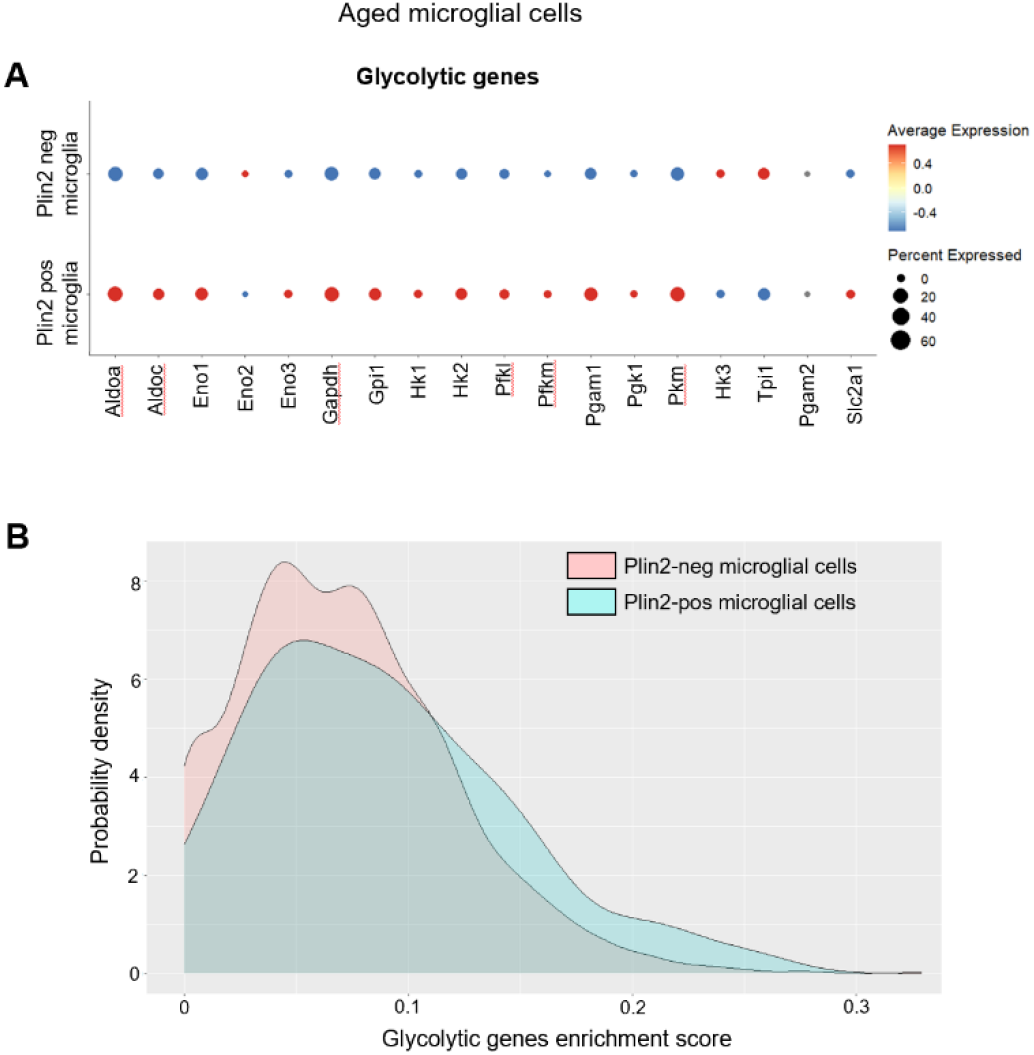
Changes in glycolytic gene expression in Plin2-expressing microglial cells. **A**) Dot plots showing the expression levels of genes involved in the glycolytic pathway in Plin2-positive and negative microglial cells from the aged brains. C) Density plots for gene set enrichment scores for glycolytic genes in Plin2-positive and negative microglia from the aged brains.

To further characterize the impact of lipid accumulation on endothelial cell function, we conducted unbiased differential gene expression analysis between Plin2-positive and Plin2-negative ECs in the aged brain. As shown in Table 2, five differentially expressed genes (DEGs) were identified—three upregulated and two downregulated in lipid-accumulating ECs. Among the upregulated genes, *Scp2* (Sterol carrier protein 2) and *Cyb5a* (Cytochrome b5) are directly involved in lipid metabolism, specifically the regulation of lipid transport between cellular compartments and lipid biosynthesis, respectively. The third upregulated gene, Ubald2, is less well characterized, and its function in ECs remains unclear. Notably, both downregulated genes, *mt-Nd4* and *mt-Co1*, encode mitochondrial electron transport chain complexes I and IV, respectively, suggesting that lipid accumulation could potentially impair mitochondrial metabolism in brain ECs.

**Table 2.** Differentially expressed genes in Plin2-positive versus Plin2-negative endothelial cells (ECs) from aged brains are shown in the table. Upregulated genes are highlighted in the first three rows with a light red shade, while downregulated genes are indicated in light blue-shaded rows.

| Genes | p_value | Fold change (avg_log2) | p_value_adjusted |
| --- | --- | --- | --- |
| Scp2 | 9.729554e-13 | 0.1849848 | 1.430147e-08 |
| Ubald2 | 1.790669e-11 | 0.4248026 | 2.632105e-07 |
| Cyb5a | 2.522719e-11 | 0.1503088 | 3.708145e-07 |
| mt-Co1 | 3.211926e-11 | -0.1856621 | 4.721210e-07 |
| mt-Nd4 | 5.731830e-11 | -0.1856486 | 8.425216e-07 |

### Lipid accumulation is associated with senescence and increased inflammation in the ECs of the aging brain

LDAMs are characterized by increased reactive oxygen species production and increased secretion of pro-inflammatory cytokines^2,24^. Similarly, lipid accumulation has been linked with cellular senescence, a stress response to macromolecular damage, in dopaminergic neurons, leading to neurodegeneration in Parkinson’s disease^25^. Given the link between lipid accumulation, inflammation, and senescence in microglia and neurons, we next investigated whether a similar phenotype is observed in ECs during aging. We performed gene enrichment score analysis for senescence core genes (n=10), senescence-associated secretory phenotype related genes (SASP, n=42), and neuroinflammation-related genes (n=89). The gene list is based on previous publications that identified senescent endothelial and microglial cells in the aging brain^26^. Our analysis revealed that Plin2-positive ECs in the aging brain displayed a rightward shift in the density plots, indicating a higher enrichment score for senescence, SASP, and inflammation-related gene sets compared to their counterparts in the young brain **(Fig. 6A-C)**. A similar increase in inflammatory gene expression was observed when comparing Plin2-positive versus negative ECs within the aging brain **(Suppl. Fig. 2)**. Parallel analysis of Plin2-positive microglial cells validated previous findings on LDAMs and showed that lipid accumulation drives an inflammatory phenotype in the microglial cells of the aging brain **(Fig. 7A-B)**. In contrast, Plin2 expression in astrocytes did not correlate with increased expression of senescence or inflammatory genes in the aging brain **(Suppl. Fig. 3)**. We also found a ∼ 2-fold increase in the percentage of senescent cells in the pooled Plin2-positive cells (6.5%) in the aging brain when compared to their counterparts in the young brain (2.7%) **(Fig. 7C)**. No changes in senescent cell population were observed in the Plin2-negative cells of the aging brain **(Fig. 7C)**. Together, these findings suggest that lipid accumulation is associated with increased senescence and inflammation, specifically in the endothelial and microglial cells during aging.

**Figure 6:**
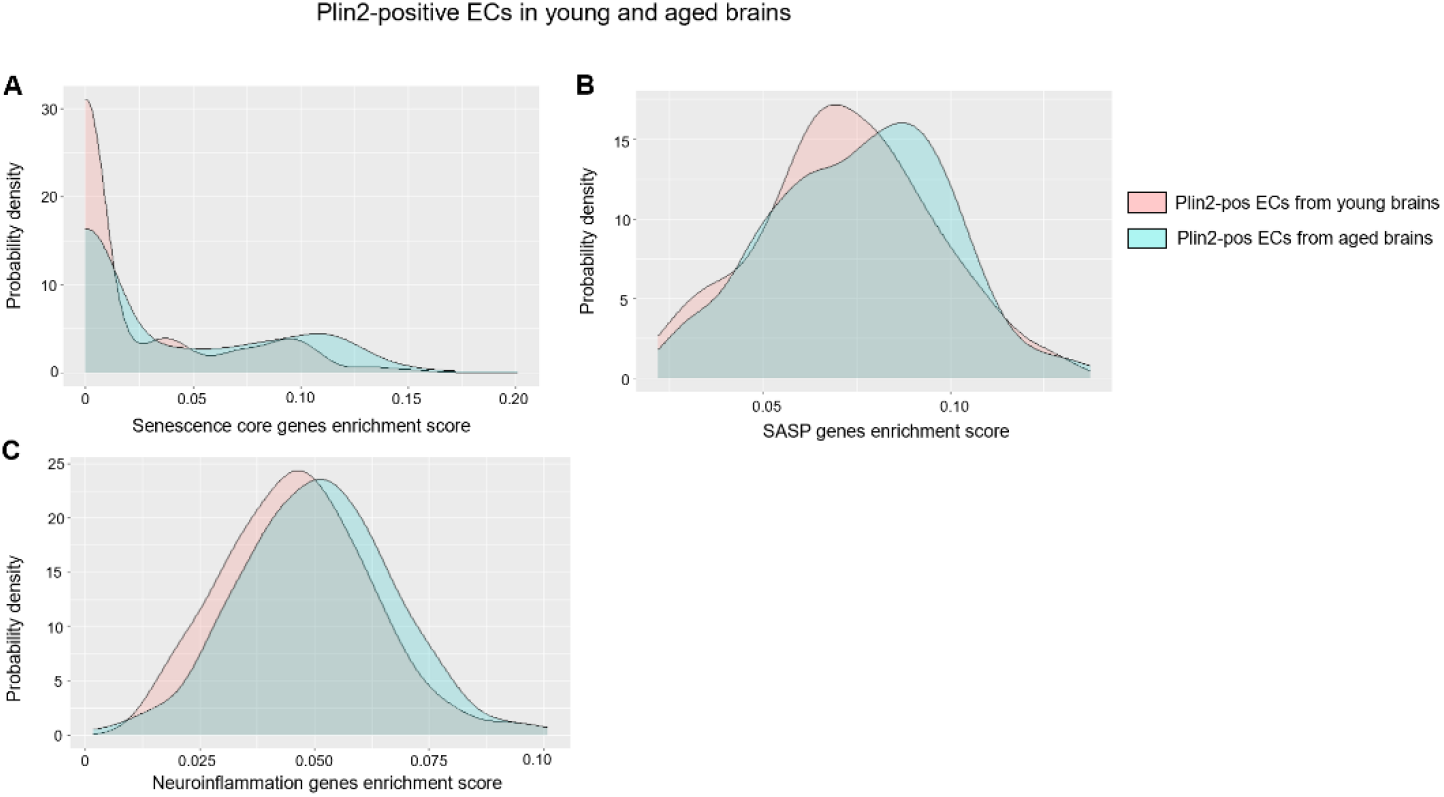
Lipid accumulation-induced changes in senescence and inflammation-related genes in young and aged brain ECs. **A-C)** Density plots depicting the gene set enrichment scores for core senescence genes, SASP- and neuroinflammation-related genes, respectively, in young and aged Plin2-expressing brain ECs.

**Figure 7:**
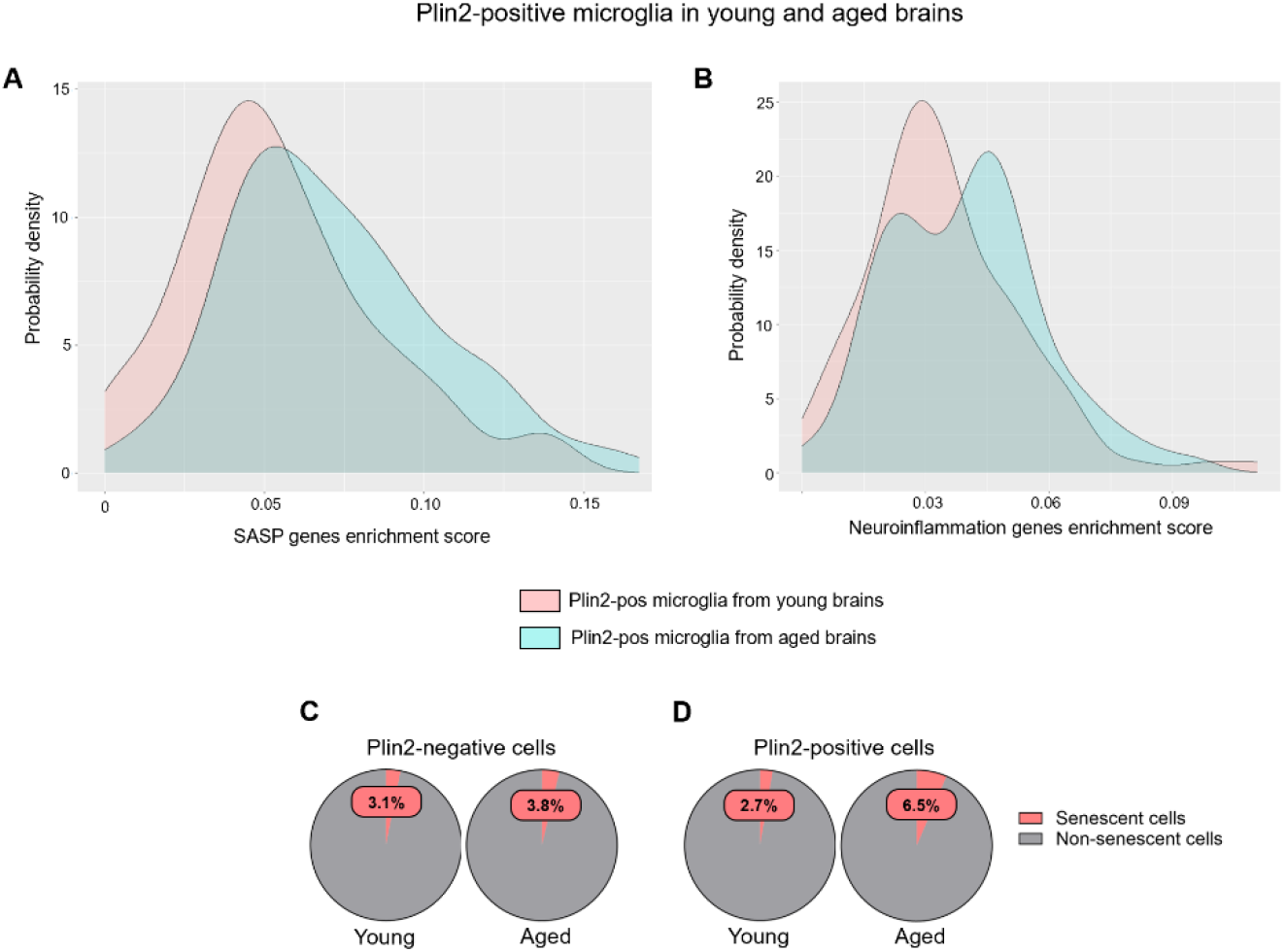
Lipid accumulation-induced changes in senescence and inflammation-related genes in young and aged brain microglial cells. **A-B)** Density plots depicting the gene set enrichment scores for SASP- and neuroinflammation-related genes, respectively, in young and aged Plin2-expressing brain microglial cells. C-D) Pie charts showing the percentage of Plin2-negative cells (panel C) and Plin2-positive cells (panel D) with high expression of senescence markers in young and aged brains.

### Validation of LD accumulating ECs in the aging brain

To corroborate our findings from sc-seq data analysis, we isolated ECs from young and aged mice brains and quantified the protein levels of Plin2, ATGL and DGAT1. In line with our transcriptomic findings, the protein levels of Plin2 were significantly higher in the CD31+ enriched cell fraction (EC fraction) from the aging brain, while it was undetectable in the young brain **(Fig. 8A-B)**. In addition, we also found that aged ECs have significantly reduced ATGL levels (reduced lipolysis) with concomitantly higher DGAT1 levels (higher lipid synthesis) **(Fig. 8A-B)**, suggesting impaired lipid turn over in the ECs with aging.

**Figure 8:**
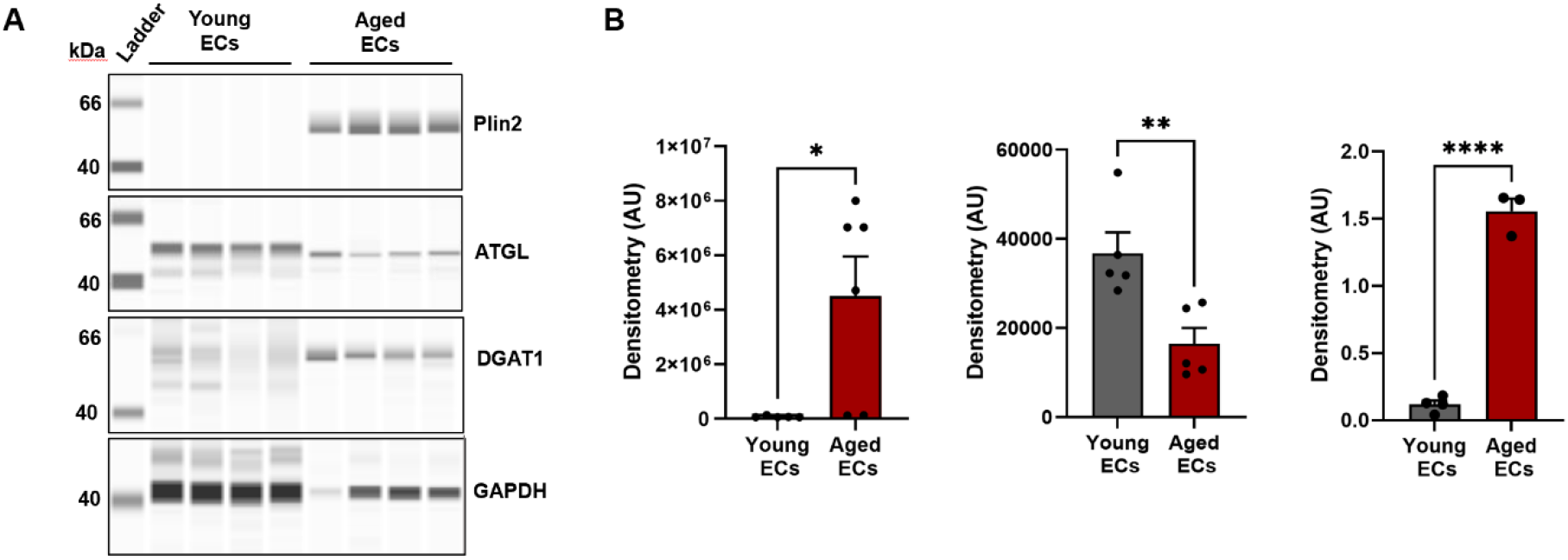
Validation of lipid accumulating ECs in the aging mouse brain. **A)** Simple Western analysis showing bands of Plin2, ATGL, DGAT1, and GAPDH protein expression in endothelial cells isolated from the brains of young and aged mice. Representative chemiluminescent images of capillary-based immunoassays showing protein bands for all the 4 protein targets. **B)** Bar graphs quantifying the protein levels after normalization to GAPDH. n=3-6/group, *p<0.05 by student’s t-test between young and aged groups.

## Discussion

In this study, we performed a secondary analysis of the single-cell transcriptomic data from young and aged mouse brain cells, with a specific focus on characterizing changes in lipid metabolism and their impact on endothelial aging. The transcriptomic signature of the aged brain ECs revealed that: 1) aging is associated with reduced lipid turn over and lipid accumulation in the brain ECs; 2) LD accumulating ECs exhibit a dysfunctional phenotype with impaired BBB integrity, increased senescence and inflammation; and 3) LD accumulation induces metabolic reprogramming in aged brain ECs, characterized by a shift from glycolysis to fatty acid oxidation and decreased mitochondrial metabolism. Together, these transcriptomic findings provide new insights into the role of dysregulated lipid metabolism in brain microvascular aging.

One of the notable findings from our study is the distinct cell-type-specific effects of lipid accumulation on metabolic and inflammatory pathways in the aging brain. In ECs, LD accumulation is associated with downregulation of key glycolytic enzymes, including *Hk1, Hk2, Pkm*, and *Slc2a1*, suggesting a potential metabolic shift in substrate preference from glucose to lipids for energy. In contrast, LDAMs exhibit upregulation of glycolytic genes, consistent with a pro-inflammatory, glycolysis-driven activation state^21-23^. Furthermore, LD accumulation is linked to increased expression of inflammatory and senescence-related genes in both ECs and microglia, but not in astrocytes, underscoring a cell-type-specific vulnerability to lipid dysregulation during aging. Although the precise mechanisms underlying this resilience in astrocytes remain unclear, their physiological role in lipid trafficking and metabolic support to neurons may have equipped them with enhanced capacity for lipid oxidation, thereby protecting against age-related lipotoxicity^1,27,28^.

The mechanisms through which lipid accumulation could induce endothelial dysfunction are multifaceted. First, our data shows that LD-accumulating ECs have increased senescence and inflammation, both are well-known players of microvascular aging^29^. Age-related oxidative stress and subsequent macromolecular damage trigger the induction of senescence, which results in irreversible proliferative arrest and induces a pro-inflammatory secretory phenotype in brain ECs^30,31^. Senescence results in pathological alterations in the structure and function of ECs, leading to BBB impairment and dysregulation in cerebral blood flow responses^30,32-35^. Studies estimate that ∼10% of ECs in the aging brain are senescent^26,32^ and eliminating senescent ECs improves BBB function, microvascular density, and cerebral blood flow responses^32,35,36^. Although senescent cells have been reported to accrue lipids^37,38^ and promote lipid accumulation in neighboring non-senescent cells^39^, whether lipid accumulation can trigger senescence in ECs remains unknown. Our results showed that the percentage of senescent cells was nearly twice as high in Plin2-positive compared to Plin2-negative cells in the aging brain, whereas no such association between Plin2 expression and senescence was observed in the young brain. These findings suggest a synergistic interaction between aging and lipid accumulation in promoting brain cell senescence.

Secondly, LD-induced metabolic reprogramming could negatively affect mitochondrial metabolism, contributing to endothelial dysfunction. Unlike astrocytes, which possess a high buffering capacity for mitochondrial oxidative stress^40^, ECs preferentially rely on glycolysis over oxidative metabolism to limit mitochondrial reactive oxygen species (ROS) generation^20^. However, our data suggest that lipid accumulation induces a metabolic shift in energy metabolism, forcing ECs to oxidize more lipids over glycolysis for energy, potentially leading to increased mitochondrial oxidative stress. This is supported by our findings showing reduced expression of the glucose transporter and key glycolytic enzymes with simultaneous upregulation of enzymes involved in fatty acid oxidation. Furthermore, the decreased expression of genes encoding mitochondrial ETC proteins reinforces the notion that lipid accumulation-driven shift to elevated fatty acid oxidation impairs mitochondrial metabolism in brain ECs.

Overall, our findings highlight a previously underappreciated role for lipid dysregulation in driving endothelial dysfunction and potentially cognitive decline in aging. Future studies using animal models that mimic age-related lipid accumulation specifically in brain ECs will help uncover the molecular mechanisms linking lipid metabolism with microvascular dysfunction. These studies have the potential to identify therapeutic targets aimed at enhancing lipid turnover to mitigate endothelial dysfunction and cognitive impairment in aging.

**Suppl. Fig. 1:** Violin plots showing expression of lipid metabolism-related genes in various brain cell types.

**Suppl. Fig. 2: Lipid accumulation-induced changes in senescence and inflammation-related genes in ECs during aging. A-C)** Density plots depicting the gene set enrichment scores for core senescence genes, SASP- and neuroinflammation-related genes, respectively, in Plin2-positive and negative brain ECs from the aged brains.

**Suppl. Fig. 3: Lipid accumulation-induced changes in senescence and inflammation-related genes in astrocytes during aging. A-C)** Density plots depicting the gene set enrichment scores for core senescence genes, SASP- and neuroinflammation-related genes, respectively in Plin2 positive and negative astrocytes from the aged brains.

## Conflict of Interest

The authors declare that they have no conflicts of interest.

## Author Contributions

Conceptualization, PB and SO; methodology, data analysis and investigation, SO, PH, TK, DN, BP IR; writing—original draft preparation, PB and SO; review and editing, SO, PH, TK, DN, BP, IR and PB; funding acquisition, PB.

## Funding

This work was supported by grants from the American Heart Association (CDA1048544 to PB), the National Institute on Aging (K01AG073613 to PB and R01HL163775 to MS), as well as the Seed grant from Presbyterian Health Foundation, Pilot grant from Harold Hamm Diabetes Center (HHDC) and OUHSC College of Medicine Alumni Association (COMAA) to PB. This research was conducted while PB was an AFAR Grant for Junior Faculty awardee, supported by the WoodNext Foundation, a component fund administered by Greater Houston Community Foundation.

## Acknowledgement

We gratefully acknowledge Ximerakis et al^17^ for publicly sharing their single-cell RNA-sequencing dataset. Their contribution made our secondary analysis and this publication possible.

## Notes

https://doi.org/10.1038/s41593-019-0491-3

### Competing Interest Statement

The authors have declared no competing interest.

