## Supplementary figures and images for "Lipid-laden endothelial cells exhibit a transcriptomic signature linked to blood-brain barrier dysfunction, metabolic reprogramming and increased inflammation in the aging brain"

### Supplemental Figure 1

#### Slide 1
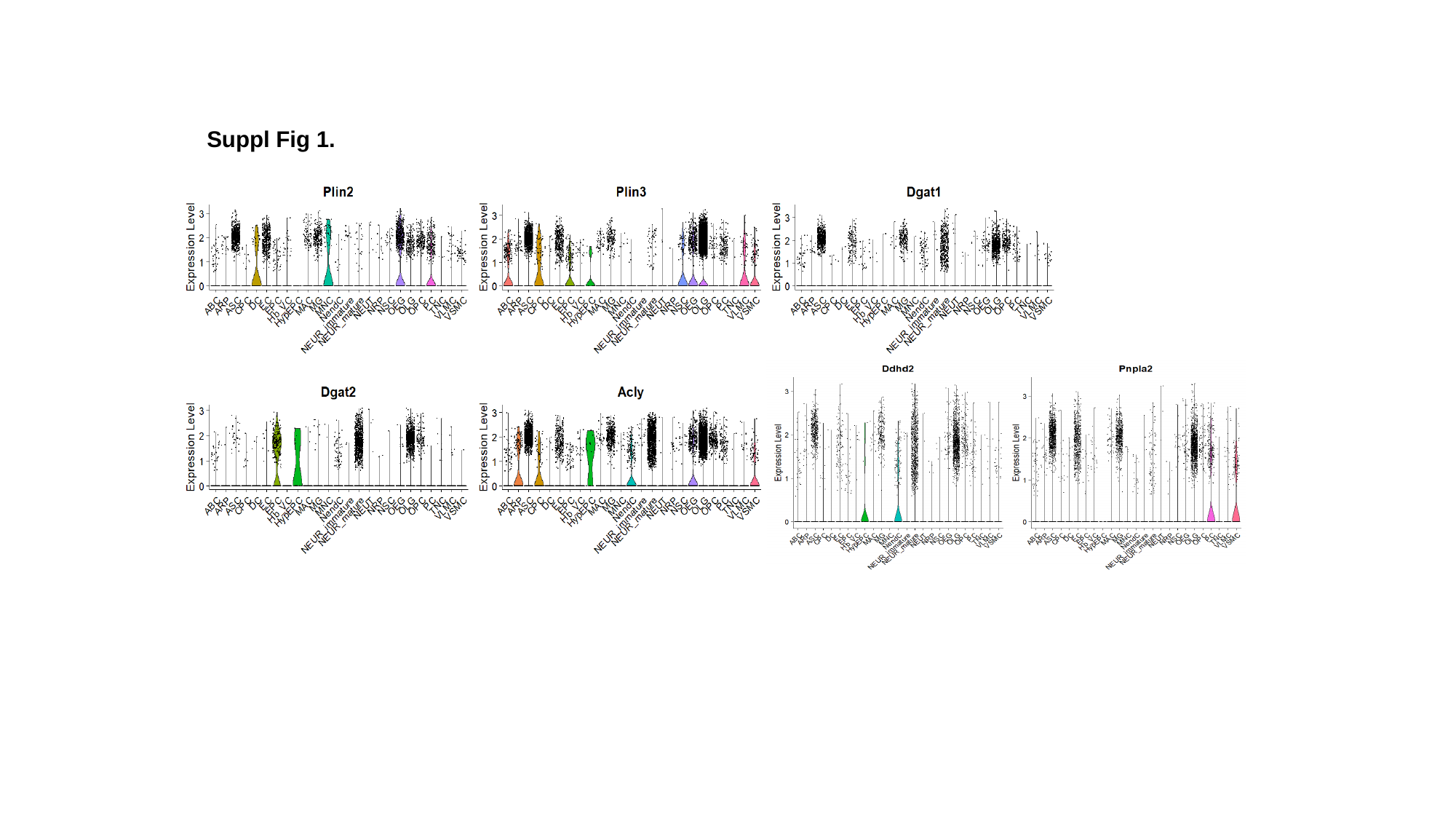

Suppl Fig 1.
