## Supplemental Figure 2 for "Lipid-laden endothelial cells exhibit a transcriptomic signature linked to blood-brain barrier dysfunction, metabolic reprogramming and increased inflammation in the aging brain"

### Slide 1
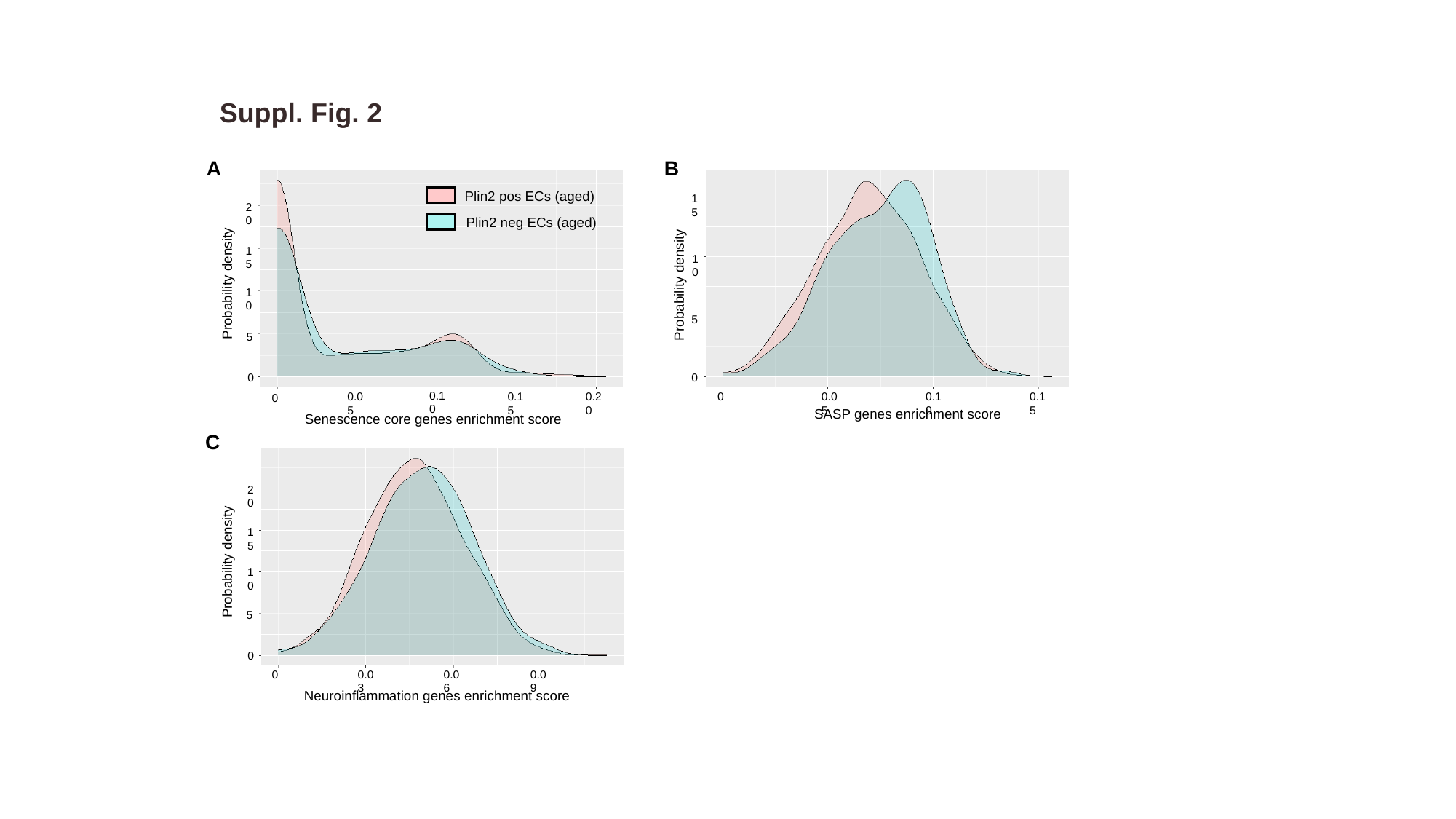

Suppl. Fig. 2
B
A
20
15
Probability density
10
5
0
0.10
0.05
0.15
0.20
0
Senescence core genes enrichment score
15
10
Probability density
5
0
0.10
0.15
0.05
0
SASP genes enrichment score
Plin2 pos ECs (aged)
Plin2 neg ECs (aged)
C
20
15
Probability density
10
5
0
0.03
0.06
0.09
0
Neuroinflammation genes enrichment score
