## Supplemental Figure 3 for "Lipid-laden endothelial cells exhibit a transcriptomic signature linked to blood-brain barrier dysfunction, metabolic reprogramming and increased inflammation in the aging brain"

### Slide 1
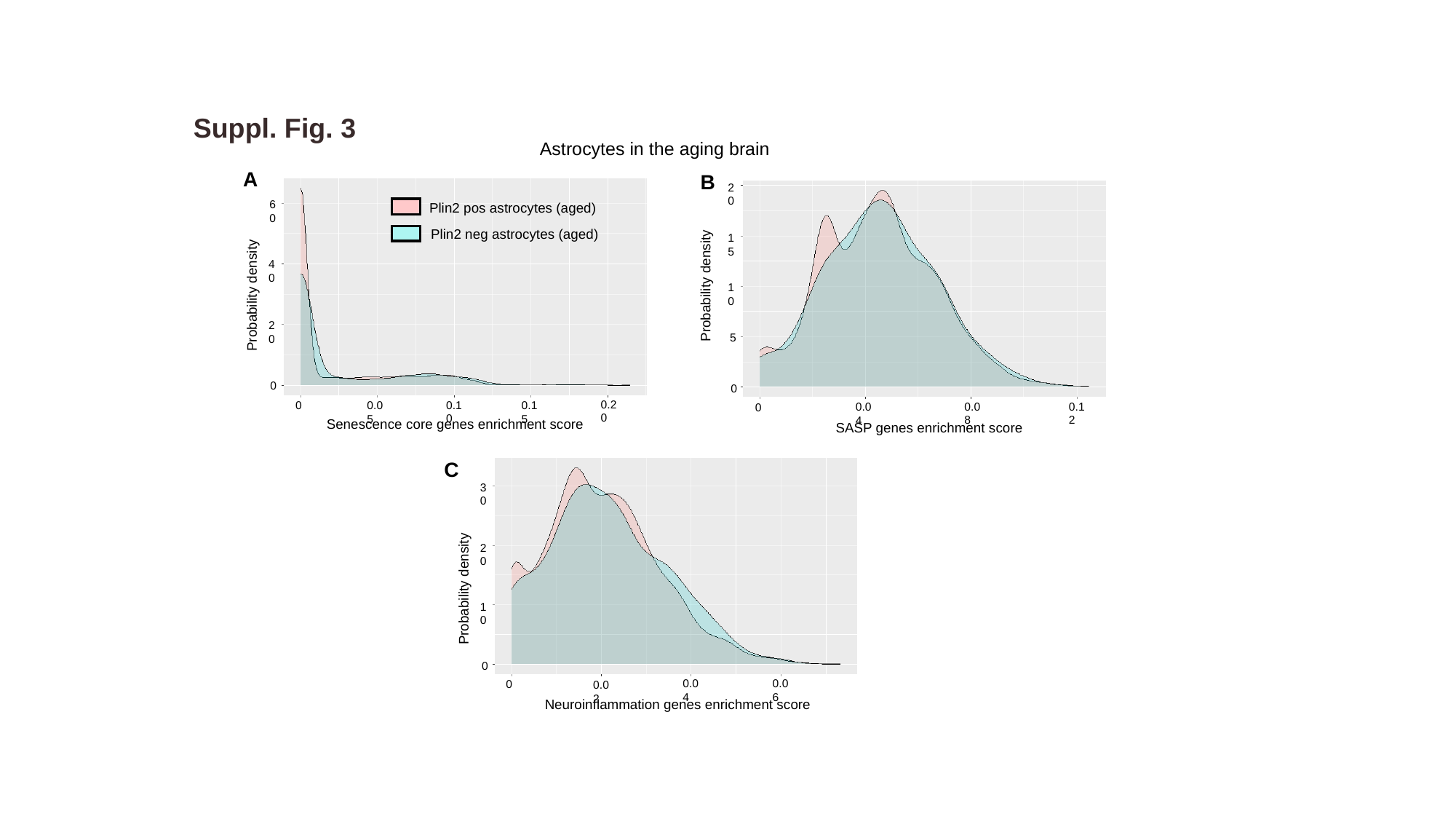

Suppl. Fig. 3
Astrocytes in the aging brain
A
B
60
40
Probability density
20
0
0.20
0.10
0.15
0
0.05
Senescence core genes enrichment score
20
15
Probability density
10
5
0
0.08
0.12
0.04
0
SASP genes enrichment score
Plin2 pos astrocytes (aged)
Plin2 neg astrocytes (aged)
C
30
20
Probability density
10
0
0.06
0.04
0
0.02
Neuroinflammation genes enrichment score
